# ATG-3 limits Orsay virus infection in *C. elegans* through regulation of collagen pathways

**DOI:** 10.1101/2025.01.13.632696

**Authors:** Gowri Kalugotla, Vivien Marmerstein, Lawrence A. Schriefer, Leran Wang, Stephanie Morrison, Luis Casorla Perez, Tim Schedl, Stephen C. Pak, Megan T. Baldridge

## Abstract

Autophagy is an essential cellular process which functions to maintain homeostasis in response to stressors such as starvation or infection. Here, we report that a subset of autophagy factors including ATG-3 play an antiviral role in Orsay virus infection of *Caenorhabditis elegans*. Orsay virus infection does not modulate autophagic flux, and re-feeding after starvation limits Orsay virus infection and blocks autophagic flux, suggesting that the role of ATG-3 in Orsay virus susceptibility is independent of its role in maintaining autophagic flux. *atg-3* mutants phenocopy *rde-1* mutants, which have a defect in RNA interference (RNAi), in susceptibility to Orsay virus infection and transcriptional response to infection. However, *atg-3* mutants do not exhibit defects in RNAi. Additionally, *atg-3* limits viral infection at a post-entry step, similar to *rde-1* mutants. Differential expression analysis using RNA sequencing revealed that antiviral *sqt-2*, which encodes a collagen trimer protein, is depleted in naïve and infected *atg-3* mutants, as well as in infected WT animals, as are numerous other collagen genes. These data suggest that ATG-3 has a role in collagen organization pathways that function in antiviral defense in *C. elegans*.

## INTRODUCTION

Autophagy is an evolutionarily-conserved process by which cells reallocate nutrients and eliminate unwanted materials such as defective proteins and damaged organelles (1). Because of its critical role in maintaining cellular homeostasis in conditions of nutrient abundance and stress, the dysregulation of autophagy is linked to many human pathologies ranging from neurodegeneration to cancer to susceptibility to pathogens (2). Indeed, autophagy has recently been implicated not only in the breakdown of phagocytosed pathogenic material but also in regulation of immune signaling pathways including interferon (IFN) signaling (2, 3). Autophagy genes and proteins have been linked to inflammation during infection as well as to innate and adaptive immune regulation (3, 4). Autophagy adaptors can also serve as their own category of pattern recognition receptors, helping to control intracellular microbes via direct capture, stimulation of inflammatory signaling, and delivery of antimicrobial peptides (4).

The steps of macroautophagy, hereafter referred to as autophagy, begin with *de novo* formation of a cup-shaped double membrane in the cytoplasm called the isolation membrane or phagophore (5). Autophagy induction begins at the phagophore assembly site, at which nucleation, the recruitment of various autophagy-related proteins, then occurs (6). After initiation, the phagophore elongates and expands, engulfing its cargo in the process (1). This proceeds to closure and completion of the structure, which is then referred to as the autophagosome (7). The autophagosome then docks and fuses with an endosome or lysosome, and the autophagic process attains completion with breakdown and degradation of the autophagosome inner membrane and cargo and recycling of the resulting macromolecules (1).

ATG3 protein is involved in the elongation of the autophagosomal membrane through its role in LC3 lipidation (8). In this process, ATG3 acts as an E2-like-conjugated enzyme and has membrane binding functions (9, 10). ATG3 is highly conserved in eukaryotes, and it is essential for homeostasis of the organism. *Atg3^-/-^* mice and *atg-3* null mutants in *C. elegans* are nonviable, requiring use of conditional or hypomorphic alleles for study (11, 12, 13). ATG3 is important in numerous biological processes, including tumor progression, maintenance of mitochondrial homeostasis, and control of infectious pathogens (14, 15, 16). ATG3 also has a role in late endosome function that is distinct from its role in autophagosome formation (17). The ATG12-ATG3 complex interacts with the endosomal sorting complexes required for transport (ESCRT)-associated protein Alix and controls several different Alix-dependent processes including late endosome distribution, exosome biogenesis, and retroviral budding (17).

Here, we demonstrate a novel role for ATG-3 in Orsay virus infection in *C. elegans*. Orsay virus is the only known virus that naturally infects *C. elegans* and was discovered from surveys of wild nematodes in Orsay, France (18, 19). Although it has yet to be formally classified, Orsay virus resembles known nodaviruses, which infect fish and insects and significantly impact the aquaculture industry (18, 20). Orsay virus is a non-enveloped, single-stranded, positive-sense RNA virus with a genome composed of two RNA segments (18, 21). The RNA1 segment, which is about 3.4 kb, encodes an RNA-dependent RNA polymerase (RdRp), and the RNA2 segment, which is about 2.5 kb, encodes the viral capsid and a capsid-delta fusion protein that is generated by a ribosomal frameshift mechanism (21).

Orsay virus is transmitted through the fecal-oral route, infects *C. elegans* intestinal cells, and causes changes in intestinal morphology (18). In addition to enlarged intestinal lumens, infected animals have subcellular structural changes such as reorganization of the terminal web, diminished intermediate filaments, and liquefaction of the cytoplasm (18, 22). In *C. elegans*, the terminal web is a subapical structure in the intestine that functions as an anchor for actin bundles that protrude from the brush border microvilli (23). The main components of the terminal web are intermediate filaments, myosin, spectrin, and several different actin-binding proteins (23, 24). The terminal web and intermediate filaments are important structural components in *C. elegans* intestinal cells. Infection with Orsay virus also causes fusion of intestinal cells, induction of vesicles of unknown type, and disappearance of nuclei (18, 19).

The discovery of Orsay virus has facilitated the use of *C. elegans* as a model system to investigate host responses to viral infection. Although *C. elegans* lack IFNs, they have a program of highly conserved innate immune response genes whose expression is induced following pathogen infection called the intracellular pathogen response (IPR), which is similar to induction of IFN-stimulated genes (25, 26). The IPR is thus named because it is activated in response to both viral infection and infection by pathogens in the microsporidian phylum, which are obligate intracellular organisms that naturally infect *C. elegans* (27, 28). One of the other main antiviral mechanisms in *C. elegans* is RNA interference (RNAi), and Dicer-related helicase (DRH-1), a homolog of mammalian RIG-I, was found to interact with the dsRNA-binding proteins RDE-1, RDE-4 and Dicer/DCR-1 to process dsRNA into small interfering RNAs (29). DRH-1 also triggers the IPR through a mechanism distinct from its role in the antiviral RNAi pathway (30). Recently, collagen genes and regulators of the actin network and the terminal web have also been identified as antiviral genes (31). Intestinally-expressed collagens may form a physical barrier along with the terminal web to prevent Orsay virus entry or egress in intestinal cells (31).

In this study, we discovered that a subset of autophagy genes including *atg-3* have an antiviral role in Orsay virus infection in *C. elegans.* We also found that re-feeding after starvation limits Orsay virus infection and blocks autophagic flux, though Orsay virus infection itself does not modulate autophagic flux. We determined that both *atg-3* disruption and Orsay virus infection altered collagen organization pathways, which have recently been identified to have an antiviral role in Orsay virus infection. Overall, our study identified a novel role for ATG-3 in structural organization of *C. elegans* cells as well as in Orsay virus pathogenesis.

## RESULTS

### A subset of autophagy proteins limit Orsay virus infection

Based on previously identified roles for autophagy in the regulation of mucosal viruses, we sought to explore whether autophagy regulates viral infection in *C. elegans* (32, 33, 34). To test this, we studied a subset of autophagy genes with critical roles in the different steps of autophagy (**Fig 1A**). Using *C. elegans* lines with mutations in these genes, we synchronized worms at the L1 stage by bleaching adults and allowing the harvested eggs to hatch overnight in M9 buffer, then allowed these worms to grow for 24h on nematode growth medium (NGM) plates seeded with OP50 *Escherichia coli*, and then orally inoculated with Orsay virus, which was added directly to the bacterial lawn.

**Figure 1:**
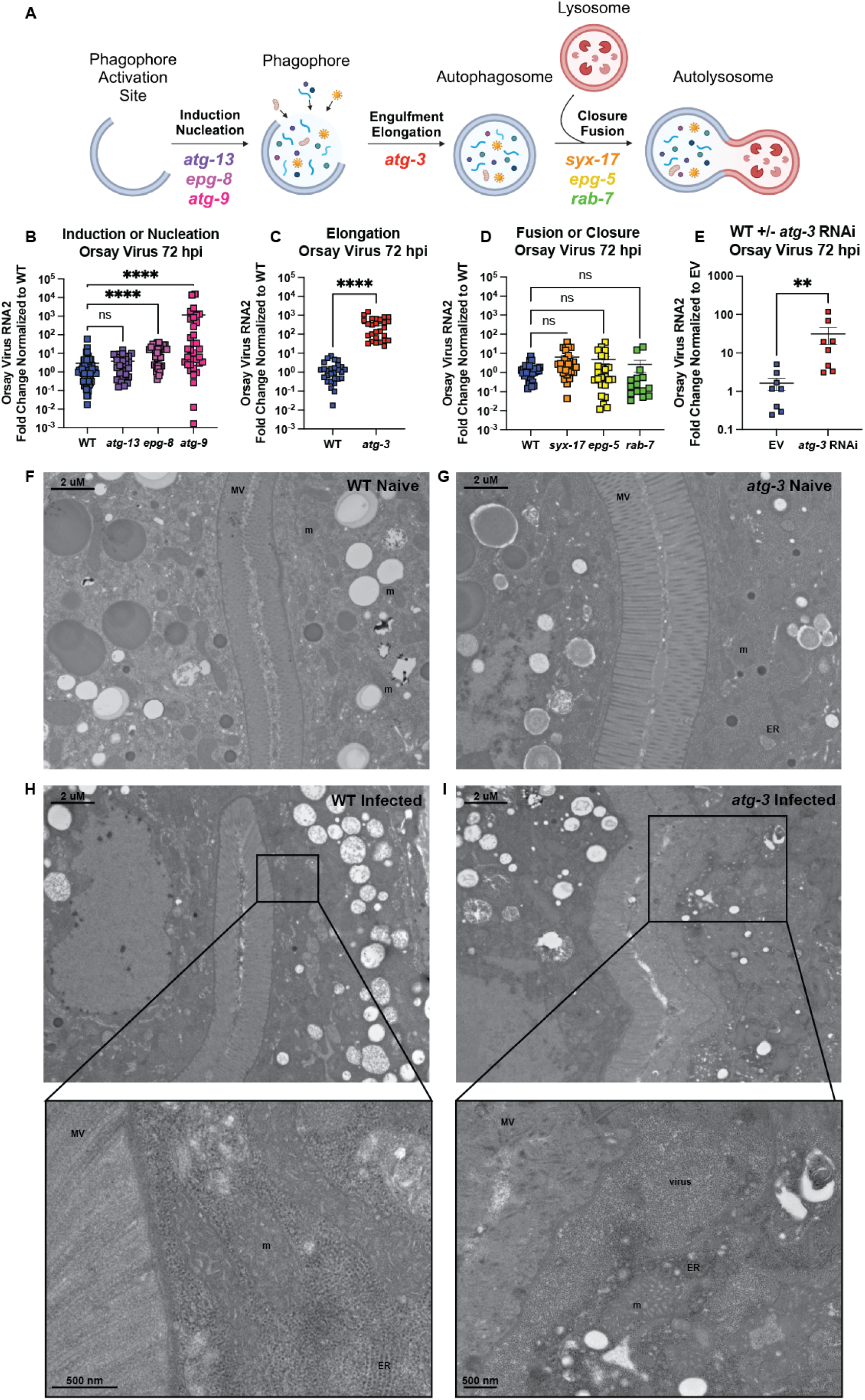
A subset of autophagy genes have an antiviral role in Orsay virus infection. **(A)** A schematic of the stages of autophagy and the selection of autophagy genes used. **(B-D)** Orsay virus infection in a subset of autophagy mutant *C. elegans* at 72 hours post infection (hpi). Wild-type (WT) and mutant animals as indicated in each graph were infected with Orsay virus, collected at 72 hours post infection (hpi), and viral RNA levels were determined with RT-qPCR. Data are from at least three independent experiments performed with six replicates. Each point represents 200 animals, and mean +/- standard error is shown. **(E)** Orsay virus infection 72 hpi in WT *C. elegans* treated with empty vector (EV) RNAi or *atg-3* RNAi. Viral RNA levels were determined with RT-qPCR. Data are from two independent experiments performed with six replicates. Each point represents 200 animals, and mean +/- standard error is shown. **(B-E)** Data analyzed by Kruskal-Wallis test with *post hoc* comparisons analyzed by Dunn’s multiple comparison test (****, p <0.0001; ***, p < 0.001, ns = non-significant p > 0.05) when three or more samples were compared. When two samples were compared, data was analyzed by Mann-Whitney test (**** = p<0.0001). **(F-I)** Transmission electron microscopy of naïve and infected WT and *atg-3* mutant *C. elegans*. These images represent four experimental replicates of 100 animals infected per sample. Scale bars show 2μM or 500 nM as labeled. Mitochondria are labeled “m,” microvilli are “MV,” the rough endoplasmic reticulum is labelled “ER,” and inclusion of viral particles is labelled “virus.”

We quantified the Orsay virus RNA2 segment by RT-qPCR 72 hours post infection (hpi). We found that mutation of *atg-13*, which encodes ATG-13 that is part of the pre-initiation complex involved in the induction of autophagy, does not change the amount of Orsay virus RNA compared to wild-type (WT) *C. elegans* (**Fig 1B**) (35). However, *epg-8* and *atg-9* mutant worms both exhibited enhanced levels of Orsay virus, suggesting antiviral roles for these factors (**Fig 1B**). EPG-8 is a member of the initiation complex that helps form the isolation membrane on the phagophore (36), while ATG-9 is involved in nucleation and is thought to promote autophagosome biogenesis by recruiting lipids to the autophagosome (37). Mutants in *atg-3*, encoding ATG-3 which is involved in engulfment and elongation, were hypersusceptible to Orsay virus with ∼220X levels of virus detected (**Fig 1C**). In contrast, worms with mutations in autophagy factors involved in closure or fusion of the autophagosome and lysosome—*syx-17*, *epg-5*, and *rab-7*— showed no differences in Orsay virus RNA levels compared to controls (**Fig 1D**) (38, 39). Taken together, this data suggests that either early stages of autophagy or autophagy-independent activities of EPG-8, ATG-9, and ATG-3 are involved in the control of Orsay virus.

To confirm the viral susceptibility phenotype that we observed in *atg-3* mutant *C. elegans*, we treated WT *C. elegans* with empty vector (EV) or *atg-3* RNAi, then infected the worms with Orsay virus, and quantified viral RNA with RT-qPCR at 72 hpi. We observed that WT *C. elegans* treated with *atg-3* RNAi were significantly more susceptible to Orsay virus infection than EV animals (**Fig 1E**). We further performed transmission electron microscopy on naïve and infected WT and *atg-3* mutants at 72 hpi. We did not observe any broad morphological differences between WT and *atg-3* mutant naïve animals (**Fig 1F, G**). In infected WT animals, we observed compromised mitochondria with disorganized cristae (labeled “m”) compared to WT naïve animals, which is a marker of cell stress and consistent with infection (**Fig 1H**). In infected *atg-3* mutants, we observed not only severely compromised mitochondria (“m”) but also damage to the microvilli (labeled “MV”) as well as large inclusions of viral particles (labeled “virus”) (**Fig 1I**). Viral particles are ∼23 nm whereas ribosomes in the rough endoplasmic reticulum (labeled ER) are ∼14 nm and are thus distinguishable in these images (**Fig 1H, I**). We were not able to visualize viral particles in infected WT samples though we observed evidence of cell stress. In contrast, we observed numerous prominent clusters of viral particles in the *atg-3* mutants, consistent with increased viral infection.

### Orsay virus infection does not modulate autophagic flux

Having found that autophagy factors regulate Orsay virus infection, we next sought to determine whether Orsay virus infection itself modulated autophagic activity or flux. To interrogate this, we used recently-generated autophagic flux reporter (AFR) animals (40). Briefly, the reporter is expressed as a fusion protein, GFP::LGG-1::mKate2. mKate2 is a red fluorescent protein that is cleaved by ATG4/ATG-4 and is not preferentially degraded by autophagic machinery, so it serves as an internal control. GFP::LGG-1 is incorporated into autophagosomal membranes and is consumed during autophagy. Since the same amount of GFP::LGG-1 and mKate2 is produced initially, autophagic flux is reported as the ratio of GFP to mKate2. Decreased autophagic flux leads to increases in this ratio and vice-versa (**Fig 2A**). We confirmed that *atg-3* mutants that express the AFR were more susceptible to Orsay virus infection than WT animals (**Fig 2B**), then quantified red and green fluorescence of animals with the AFR. We observed a higher GFP/mKate2 ratio in the *atg-3* mutants compared to WT animals, an expected observation given the impairment of autophagic flux in *atg-3* mutants (**Fig 2C**) (40). However, we found no differences in the GFP/mKate2 ratios between naïve and infected WT animals or *atg-3* mutants with the AFR (**Fig 2C**), indicating that autophagic flux is not altered in the context of Orsay virus infection.

**Figure 2:**
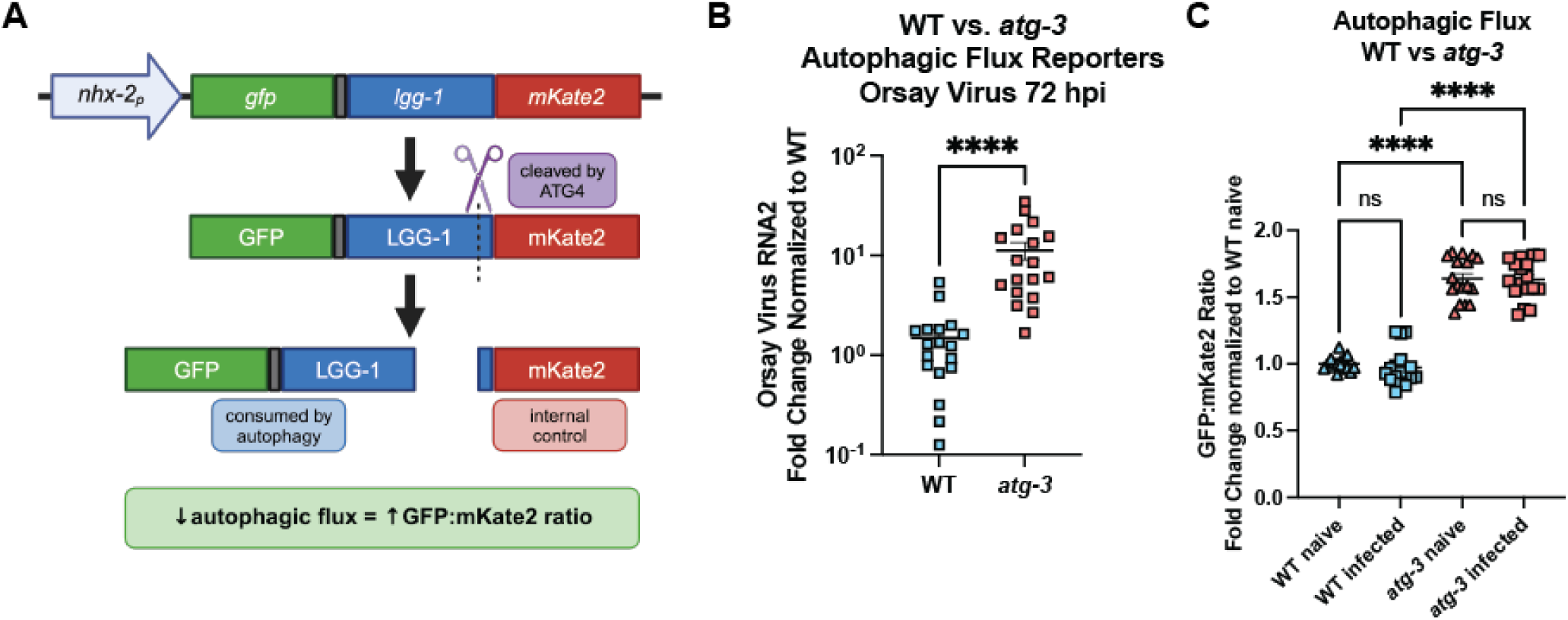
Orsay virus infection does not modulate autophagic flux. **(A)** Schematic showing how the autophagic flux reporter (AFR) is processed. The reporter is expressed in the intestine as a fusion protein, GFP::LGG-1::mKate-2, which is cleaved by ATG4/ATG-4. GFP::LGG-1 is consumed by autophagy, so decreased autophagic flux leads to accumulation of GFP::LGG-1. Cleaved mKate2 remains in the cytosol and acts as the internal control. **(B)** Orsay virus infection in adult WT and *atg-3* mutant worms expressing the AFR, 72 hpi. Viral RNA levels were determined with RT-qPCR. Data are from at least three independent experiments performed with six replicates. Each point represents 200 animals, and mean +/- standard error is shown. Statistical significance was determined by Mann-Whitney test (**** = p<0.0001). **(C)** GFP:mKate2 ratios of WT and *atg-3*-mutant animals, naïve or infected with Orsay virus 48 hpi. All data were normalized to WT naïve. Data are from 3 independent experiments, and each square represents about 80 animals. Statistically significant differences were determined by Kruskal-Wallis test with statistical difference identified between *post hoc* comparisons analyzed by Dunn’s multiple comparison test (****, p <0.0001; ns = non-significant p > 0.05).

### Re-feeding after starvation limits Orsay virus infection and blocks autophagic flux

Starvation is well-established to affect autophagy in *C. elegans* (41). Because of this, *C. elegans* with mutations in autophagy genes are passaged frequently to avoid the induction of autophagy and epigenetic changes caused by starvation. Thus, animals with the AFR cannot be starved and refed since this affects the GFP signal. While the standard Orsay virus infection protocol includes an overnight starvation step to synchronize the worms, we developed a distinct infection protocol for the AFR reporter animals to synchronize worms without overnight starvation (**Fig 3A**). We speculated that distinct infection protocols could modulate infection levels due to changes in autophagic flux. When we directly compared infection protocols, we found that unstarved animals with or without the AFR were more susceptible to Orsay virus infection than *C. elegans* synchronized with the standard infection protocol, labelled “re-fed”, by RT-qPCR 48 hpi (**Fig 3B**). Since starvation induces autophagy and mutation of some autophagy genes increased susceptibility to Orsay virus infection (**Fig 1**), we hypothesized that re-fed worms would have increased autophagic flux compared to unstarved worms. Surprisingly, we found re-fed animals had decreased flux compared to unstarved animals (**Fig 3C**). We once again did not observe any differences in autophagic flux between naïve and infected worms regardless of their starvation status (**Fig 3C**). The finding that unstarved worms were more susceptible to infection with Orsay virus than re-fed worms but have a higher level of autophagic flux suggests that the role of ATG-3 in resistance to Orsay virus infection could be independent of its role in canonical autophagy.

**Figure 3:**
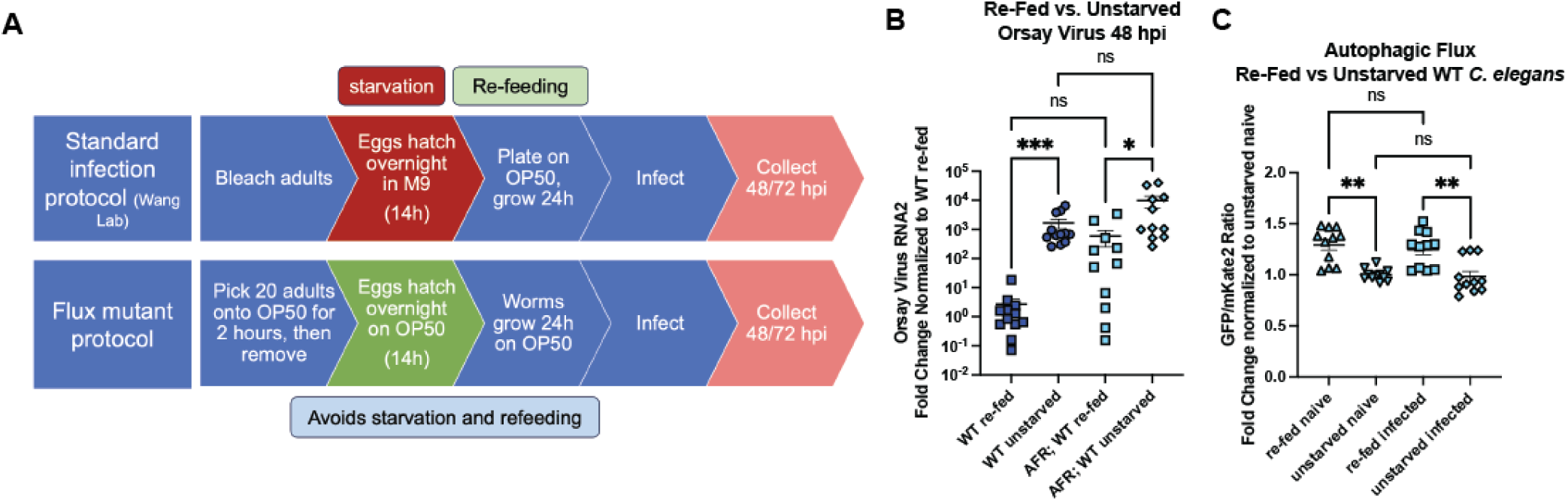
Re-feeding after starvation limits Orsay virus infection and blocks autophagic flux. **(A)** Schematic of the standard protocol for *C. elegans* synchronization and infection, which includes an overnight starvation step, as well as the modified protocol for AFR worms that does not include a starvation step. Animals were infected at 38h of life at the L3/L4 stage of development and harvested as adults. **(B)** Orsay virus infection in adult re-fed vs. unstarved WT *C. elegans* and WT *C. elegans* with the AFR, 48 hpi. Viral RNA levels were determined with RT-qPCR. Data are from two independent experiments performed with six replicates. Each point represents 200 animals, and mean +/- standard error is shown. **(C)** GFP:mKate2 ratios of re-fed vs. unstarved WT *C. elegans* carrying the AFR, naïve or infected with Orsay virus 48 hpi. All data were normalized to the unstarved naïve group. Data are from 4 independent experiments, and each square represents about 80 animals. **(B, C)** Statistical significance was determined by Kruskal-Wallis test with statistical difference identified between *post hoc* comparisons analyzed by Dunn’s multiple comparison test (****, p < 0.0001; ***, p = 0.0002; *, p = 0.0102; ns = non-significant p > 0.05).

### *atg-3*-mutant *C. elegans* phenocopy *rde-1* mutants in IPR response to infection

We next investigated whether the phenotype we observed in *atg-3* mutants was similar to that of *rde-1* mutants, which have a defect in RNAi and are known to be hypersusceptible to Orsay virus infection. We found that *atg-3* and *rde-1* mutants have similar levels of Orsay virus RNA at 72 hpi (**Fig 4A**). Since WT *C. elegans* are thought to be resistant to Orsay virus compared to the wild isolates of *C. elegans* in which Orsay virus was first isolated, much of the work to characterize the IPR has been done in *C. elegans* strains that have mutations in RNAi genes including *rde-1* (31, 42). To evaluate whether the hypersusceptibility of *atg-3* mutant worms was due to defects in antiviral responses, we evaluated induction of a commonly used panel of IPR genes between WT, *rde-1*, and *atg-3* mutants (26, 43). We analyzed IPR genes of unknown function, *F26F2.1*, *F26F2.3*, *F26F2.4*, and *pals-5* as well as four IPR genes involved in the unfolded protein response, *cul-6*, *skr-3*, *skr-4*, and *skr-5*. We found that, like *rde-1* mutants, *atg-3* mutants exhibited robust induction of IPR gene expression upon infection (**Fig 4B-I**). Further, both mutants generally exhibited enhanced infection-induced IPR gene expression compared to WT worms, likely mounting enhanced responses due to greater viral levels (**Fig 4B-I**). These data suggest that *atg-3* worms do not have a defect in the IPR response. Because we observed that *atg-3* worms phenocopied *rde-1* worms, we generated *atg-3; rde-1* double mutants to evaluate whether viral infection could be further enhanced when both genes were disrupted. Instead, we found that the double mutants had the same amount of Orsay virus RNA as *atg-3* and *rde-1* single mutants, suggesting either that the hypersusceptibility of *atg-3* and *rde-1*-mutant *C. elegans* to Orsay virus is mediated through a common pathway, or that Orsay virus accumulates at its maximal potential in each of the mutant strains and the accumulation is limited by a different factor, such as speed of replication (**Fig 4J**).

**Figure 4:**
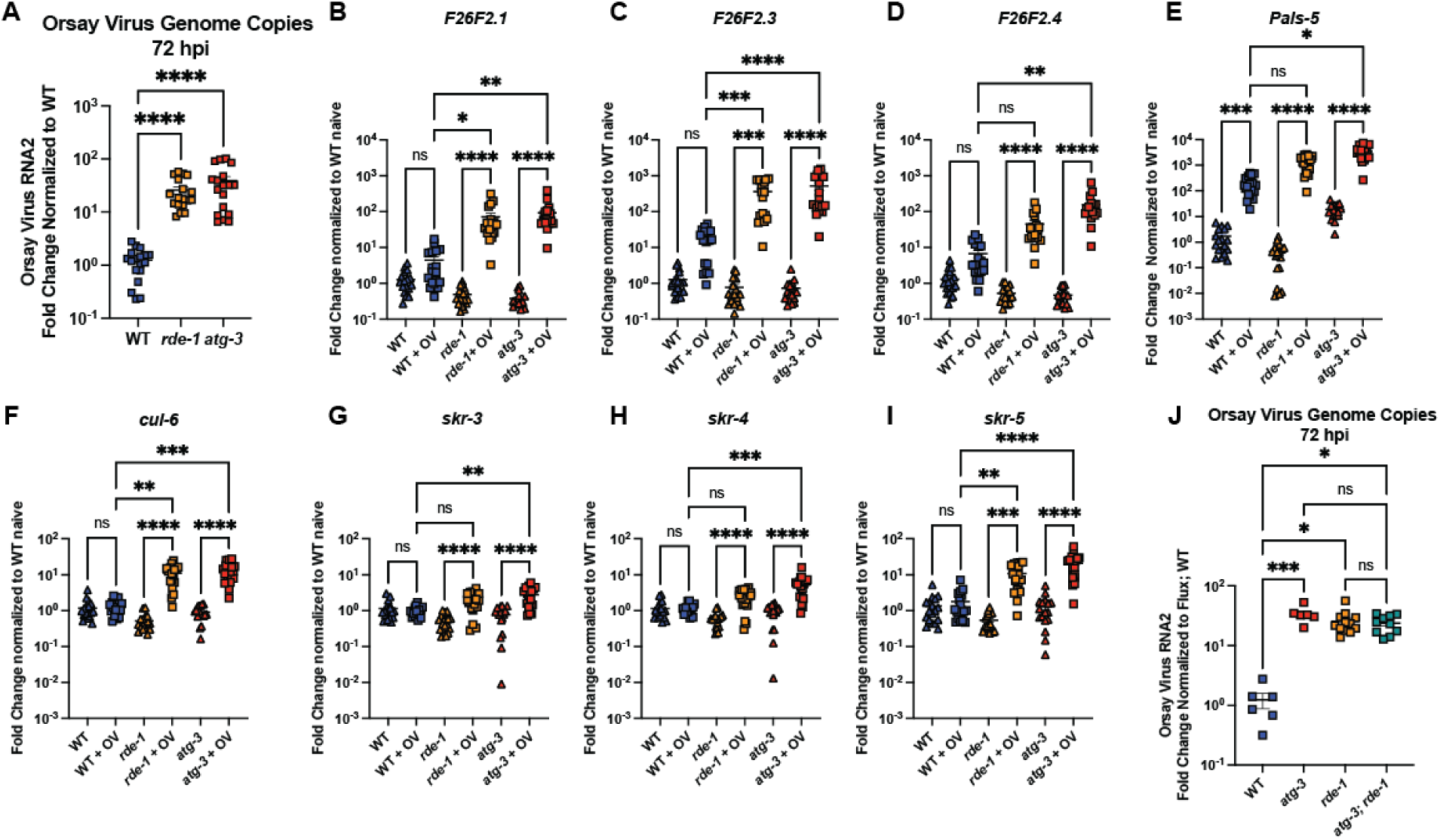
*atg-3*-mutant *C. elegans* phenocopy *rde-1* mutants in IPR response to infection. **(A)** Orsay virus infection in WT, *rde-1* and *atg-3* mutant worms, 72 hpi. Viral RNA levels were determined with RT-qPCR. Data are from 3 independent experiments performed with 6 replicates. Each point represents 200 animals, and mean +/- standard error is shown. **(B-I)** RT-qPCR measurement of IPR gene expression in WT, *rde-1*, and *atg-3* naïve and infected animals (showed as +OV), 72 hpi, shown as fold change relative to WT naïve. Data are from 3 independent experiments performed with 6 replicates. Each point represents 200 animals, and mean +/- standard error is shown. **(J)** Orsay virus infection in WT, *rde-1*, *atg-3*, and *atg-3; rde-1* mutant *C. elegans*, 72 hpi. Data are from 2 independent experiments performed with at least 3 replicates. Each point represents 200 animals, and mean +/- standard error is shown. **(A-J)** Statistical significance was determined by Kruskal-Wallis test with statistical difference identified between *post hoc* comparisons analyzed by Dunn’s multiple comparison test (****, p < 0.0001; ***, p < 0.001; **, p < 0.01; *, p < 0.05; ns = non-significant p > 0.05).

### ATG-3 mediates antiviral effects post-entry

To determine if ATG-3 was critical to limit viral levels at a pre- or post-entry step of replication, we used a previously-described replicon system for Orsay virus that is based on an extrachromosomal array of plasmids of the Orsay virus WT RNA1 segment that encodes the RdRp under a heat shock promoter (N2; PHIP::PNA1WT) (44, 45, 46). This strain is competent to support replication of the Orsay virus RNA1 segment after heat shock induction, effectively bypassing the requirement for cellular entry. As a negative control, we also used a strain carrying extrachromosomal arrays of a D601A polymerase-dead mutant of Orsay virus RNA1 (N2; PHIP::RNA1D601A). The replicon strains carry the *jyIs8* reporter, an integrated GFP reporter driven by the *pals-5* promoter that enables visualization of viral infection by fluorescence microscopy (44, 47). Using worm strains derived from crossing the replicon reporter strains to WT, *rde-1*, or *atg-3* mutant strains, we imaged worms carrying the WT and D601A RNA1 segment (**Fig 5A-F**) and counted animals that expressed *pals-*5::GFP in the intestine (**Fig 5G**). In both *rde-1* and *atg-3* mutants, significantly higher proportions of worms with the WT RNA1 segment expressed GFP, consistent with dysregulated viral replication in these lines **(Fig 5A-C, G)**. As expected, no GFP positivity was detected for any strain carrying the D601A RNA1 segment (**Fig 5D-F, G**). Thus, ATG-3 limits viral infection at a post-entry step, similar to RDE-1.

**Figure 5:**
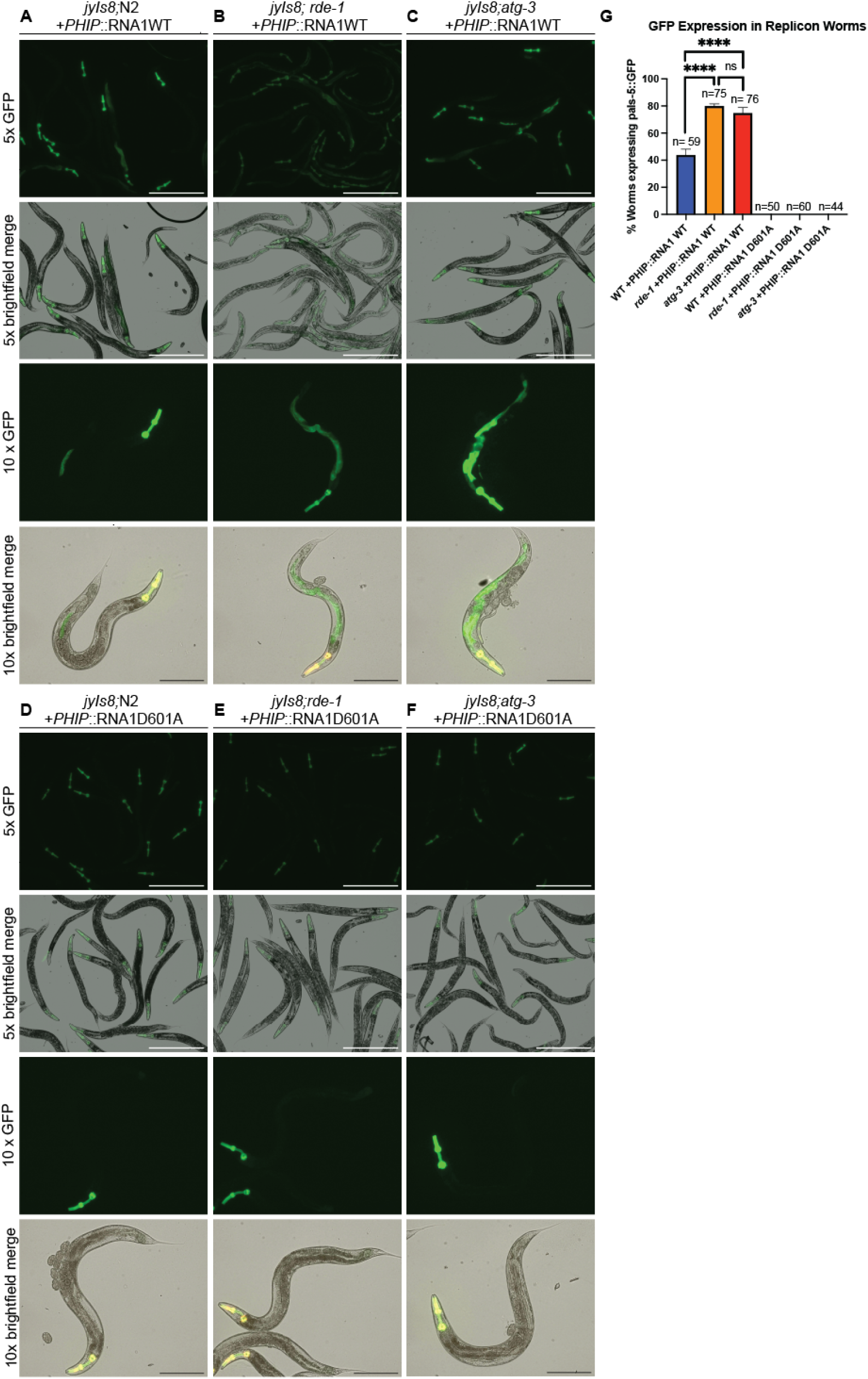
*atg-3* mutation promotes viral infection at a post-entry step. **(A-F)** Images of replicon animals of each genotype 24 hours after heatshock induction at 5x and 10x magnification. White scale bars show 500 μM, black scale bars show 250 μM. **(G)** Bar graph showing the percentage of WT, *rde-1*, and *atg-3* animals carrying the WT or D601A mutant RNA1 reporter expressing the pals-5::GFP reporter 24 hours after heatshock induction. Worms were counted as having a positive signal if the full body of the worm and the GFP reporter in the head were visible along with GFP signal in the intestine. Data are from 2 independent experiments performed with three replicates, and mean +/- standard error is shown. Total n per genotype is displayed over each bar. Statistical significance was calculated with Fisher’s Exact test. Stars are not shown for the comparisons between WT and D601A RNA1 segments, but each had p <0.0001 (****, p < 0.0001; ns = non-significant p > 0.05).

### Neither a defect in RNAi nor a defect in neutral lipid accumulation cause susceptibility to Orsay virus in *atg-3* mutant *C. elegans*

The observations that *atg-3* mutants *C. elegans* phenocopy *rde-1* mutants in the transcriptional response to infection as well as in post-entry regulation, and that *atg-3; rde-1* double mutants did not exhibit any additional enhanced viral replication (**Fig 4, 5**) led us to question whether *atg-3* mutant *C. elegans* could have an RNAi defect. We used RNAi knockdown of *pos-1*, which is required in the maternal germline for embryonic viability, and *unc-22*, which is orthologous to human *TTN* (titin) and induces muscle twitching upon RNAi knockdown (29, 48), to interrogate whether *atg-3* mutants had disrupted RNAi activity. We found that *pos-1-*knockdown led to WT worms that laid 99.9% dead embryos (n=3366) compared to 1.9% dead embryos from *rde-1* worms (n=1127) (**Fig 6A**). In *atg-3* mutants, we observed 88.3% dead embryos (n=2556), suggesting intact RNAi activity (**Fig 6A**). Similarly, upon *unc-22* RNAi knockdown, we observed that 100% of the WT and *atg-3* mutant *C. elegans* (n=283 and n=163 respectively) were twitching, compared to 0% of the *rde-1* mutants (n=402) (**Fig 6B**). Combined, these results indicate that the susceptibility of *atg-3* mutant *C. elegans* to Orsay virus is not due to a defect in RNAi function.

**Figure 6:**
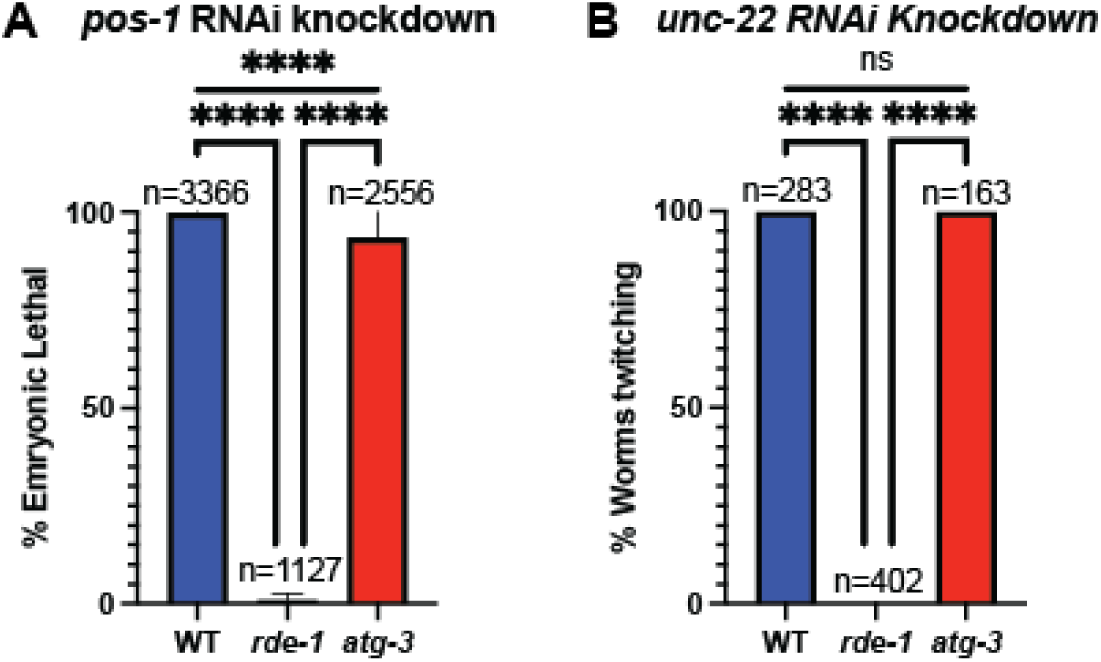
*atg-3-*mutant *C. elegans* do not have a defect in RNAi. **(A)** Bar graph showing the percent embryonic lethality in WT, *rde-1*, and *atg-3*-mutant *C. elegans*. N= number of embryos assayed in 2 separate experiments with 3 replicates each. Mean +/- standard error is shown. Animals were fed with RNAi bacteria targeting *pos-1*, which leads to progeny embryonic lethality in animals with functional RNAi. **(B)** Bar graph showing the percent of twitching animals in WT, *rde-1*, and *atg-3*-mutant *C. elegans*. N= number of embryos assayed in 2 separate experiments with 3 replicates each. Mean +/- standard error is shown. Animals were fed with RNAi bacteria targeting *unc-22*, which leads to twitching motility animals with functional RNAi. Statistical significance was determined by Fisher’s Exact test (****, p < 0.0001; ns = non-significant p > 0.05).

We next hypothesized that neutral lipid accumulation could play a role in *atg-3* mutant susceptibility to Orsay virus infection since lipid abundance can affect Orsay virus infection and vice versa (47). Mutations in autophagy genes are known to affect lipid levels in *C. elegans*, so we evaluated whether viral susceptibility in autophagy mutants correlated with neutral lipid accumulation by LipidTox staining (49). We imaged animals that had LipidTox dye mixed into their OP50 and quantified fluorescence (**Supp Fig 1A-I**). We also used *rict-1* mutants, which have increased autophagic flux and neutral lipid accumulation, as a control (**Supp Fig 1J, K**) (50). We found that the patterns of lipid accumulation did not correlate with changes in susceptibility to Orsay virus infection, suggesting this does not underlie the phenotypes observed in *atg-3* worms (r=0.1167, p=0.7759) (**Supp Fig 1L**).

**Supplemental Figure 1:**
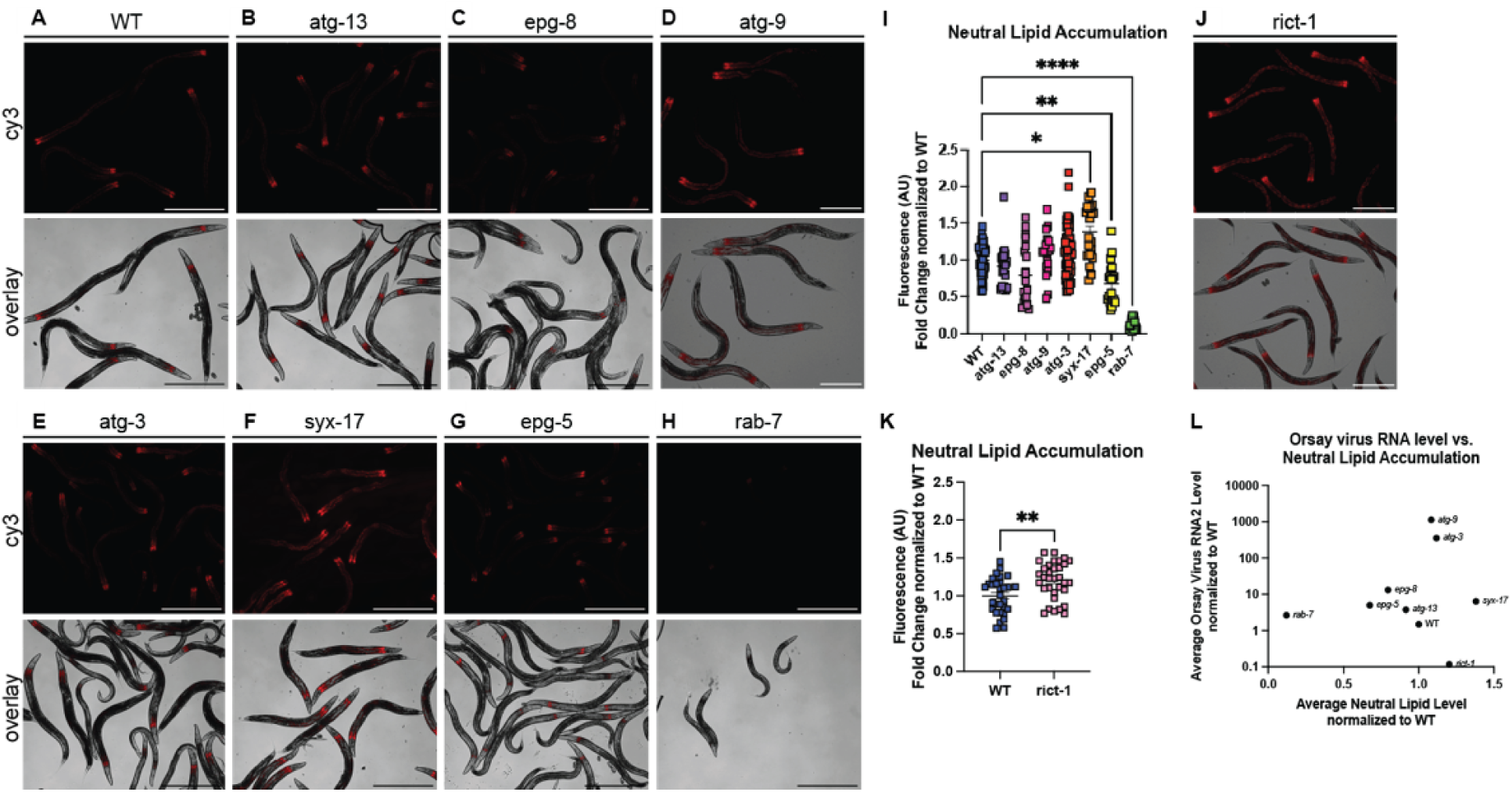
Neutral lipid accumulation in a subset of autophagy mutants does not correlate with susceptibility to Orsay virus infection. **(A-H, J)** Neutral lipid fluorescence microscopy was performed on day 3 adults that were synchronized as L1 embryos and plated on OP50 mixed with LipidTox dye. Scale bars represent 500 μM. **(I, K)** Quantification of neutral lipid fluorescence microscopy shown in panels A-H and J. Fluorescence (shown in arbitrary units (AU)) was normalized by setting the value of WT to 1. Each point represents one animal. Data are from 3 independent experiments performed with 3 replicates. Mean +/- standard error is shown. Statistically significant differences were determined by Kruskal-Wallis test with *post hoc* comparisons analyzed by Dunn’s multiple comparison test (****, p <0.0001; ***, p < 0.001; *, p <0.05; ns = non-significant p > 0.05) when three or more samples were compared. When two samples were compared, statistical significance was determined by Mann-Whitney test (** = p<0.01). **(L)** Average Orsay virus RNA levels plotted against average neutral lipid accumulation. Non-parametric Spearman correlation coefficient r=0.1167 with a p value of 0.7759.

### *atg-3* disruption affects collagen organization pathways

To investigate the pathways by which *atg-3* affects Orsay virus infection in an unbiased fashion, we performed RNAseq and differential expression analysis using DESeq2 on naïve and infected WT and *atg-3* mutant *C. elegans* at 72 hpi. We identified numerous differentially expressed genes in pairwise comparisons between these four groups (**Fig 7A-D, Supplemental Table 1**). Several of the IPR genes that we assayed by RT-qCR were also identified by our RNAseq pipeline in infected vs. naïve samples in both genotypes, including *F26F2.1*, *F26F2.3, F26F2.4,* and *skr-4* (**Fig 7A, B**) (**Fig 4B-D, H**). We also observed that *F26F2.1* and *F26F2.3* expression was increased in *atg-3* infected *C. elegans* compared to WT samples, which is consistent with our qPCR data (**Fig 7D, Fig 4B, C**).

**Figure 7:**
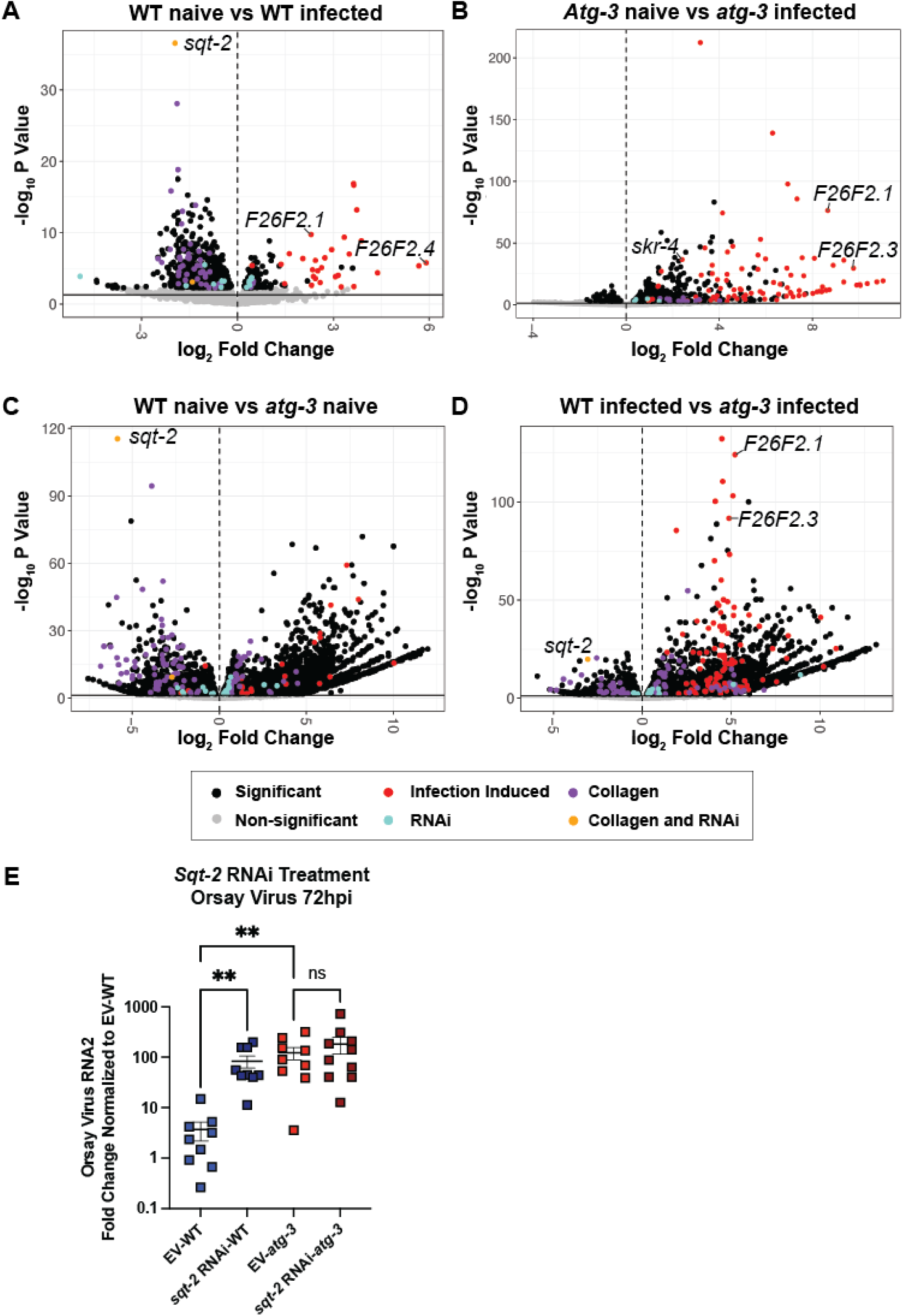
RNAseq analysis reveals that ATG-3 maintains expression of antiviral collagen genes such as *sqt-2*. RNAseq and differential expression analysis was performed on three replicates of naïve and infected WT and *atg-3*-mutant *C. elegans* 72 hpi with package DESeq2 (v. 1.34.0) to determine differences between **(A)** WT naïve vs. WT infected, **(B)** *atg-3* naïve vs. *atg-3* infected, **(C)** WT naïve vs. *atg-3 naïve*, and **(D)** WT infected vs. *atg-3* infected animals. Results were filtered for only those genes with Benjamini-Hochberg false-discovery rate adjusted p-values less than or equal to 0.05, and volcano plots were generated. Points labelled in red overlap with previously identified genes that are transcriptionally activated by Orsay virus infection at 12 hpi, and points labeled in blue were genes identified to be important for viral infection in an RNAi knockdown screen (31, 42). Points labelled in purple are genes previously identified as collagen genes, and points labelled in yellow are collagen genes that overlap with genes identified as important for viral infection in the RNAi knockdown screen (31, 51). **(E)** Orsay virus infection in WT *atg-3* mutant worms fed either empty vector (EV) or *sqt-2* RNAi, 72 hpi. Viral RNA levels were determined with RT-qPCR. Data are from 2 independent experiments performed with 6 replicates. Each point represents 200 animals, and mean +/- standard error is shown. Statistical significance was determined by Kruskal-Wallis test with statistical difference identified between *post hoc* comparisons analyzed by Dunn’s multiple comparison test (**, p < 0.01; ns = non-significant p > 0.05).

We performed pathway analysis to assess for enriched pathways associated with these differentially-expressed genes, but did not observe any pathways with significant differences emerge. We thus sought to compare our results to previous studies identifying genes or transcripts associated with Orsay virus infection. We first compared the results of our analysis to previously-published RNAseq results from WT *C. elegans* naïve versus Orsay virus-infected worms collected at 12 hpi (42). Factors previously found to be induced upon infection that also emerged as differentially-expressed in our analysis are shown in red **(Supplemental Table 2,** “Infection-induced”). We observed significant induction of transcripts consistent with the prior study upon Orsay virus infection in animals of both genotypes (WT infected vs WT naïve, p-adj=6.17×10^-23^; *atg-3* infected vs *atg-3* naïve, p-adj= 3.29×10^-43^; *atg-3* infected vs WT infected, p-adj=9.82×10^-27^)(**Fig 7A, B**). We also compared our differentially-expressed genes to a list of 108 genes identified as antiviral factors in an RNAi screen, shown in blue (**Fig 7**) (**Supplemental Table 2**, “RNAi”) (31, 42). While there was no significant pathway enrichment or depletion of these “RNAi” genes in any of our comparisons, we did identify *sqt-2* mRNA among these factors as an established antiviral gene that was one of the most significantly depleted factors in infected WT animals compared to naïve animals, naive *atg-3* mutants compared to naïve WT animals, and WT infected animals compared to *atg-3* infected animals (**Fig 7A, C, D**). *sqt-2* mRNA was not significantly different between *atg-3* naïve and *atg-3* infected samples (**Fig 7B**).

SQT-2 is predicted to be part of a collagen trimer located in membranes and is expressed in intestinal cells (31). To investigate the relationship between *sqt-2* and Orsay virus infection in *atg-3* mutants, we treated WT and *atg-3* mutant *C. elegans* with *sqt-2* RNAi vector. Consistent with the prior report, we found that WT worms fed *sqt-2* RNAi were significantly more susceptible to Orsay virus than worms treated with empty vector (EV) (31). However, treatment of *atg-3* mutant *C. elegans* with *sqt-2* RNAi did not enhance viral susceptibility (**Fig 7E**). This suggests that *atg-3* plays a role upstream of *sqt-2* activity, which modulates Orsay virus infection.

In addition to *sqt-2*, we observed mRNA depletion for several other collagen-encoding genes in naïve *atg-3* mutant *C. elegans* compared to WT animals, including *col-90*, *rol-8*, *sqt-1*, *col-17*, *col-41*, *col-107*, *dpy-5*, and *col-34* (**Supplemental Table 1**). To further explore whether collagen organization pathways are disrupted in naïve and infected WT and *atg-3* mutant *C. elegans*, we compared our differentially expressed genes to a previously published list of all collagen genes in *C. elegans* shown in purple (**Fig 7A-D**) (**Supplemental Table 2**, “Collagen”) (51), and found that this pathway was depleted in both WT infected animals compared to naïve (p-adj=2.85×10^-18^) and in *atg-3* naïve animals compared to WT naïve (p-adj=5.25×10^-12^), but was enhanced in *atg-3* infected worms compared to *atg-3* naïve (p-adj=2.27×10^-07^). Overlapping genes between the collagen gene list and antiviral genes identified by RNAi screening, including *sqt-2*, are shown in yellow (**Fig 7A-D**) (31, 51). Our data indicates broad changes in collagen organization pathways, some of which have been previously identified as antiviral factors in Orsay virus infection, in *atg-3* mutant *C. elegans* (31). The dysregulation of these pathways thus potentially leads to enhanced viral fitness in *atg-3* mutants.

## DISCUSSION

Autophagy is an essential degradative process which functions to maintain cellular homeostasis and respond to external stressors such as starvation (52). Here, we report that autophagy factors ATG-3, ATG-9, and EPG-8 play an antiviral role in Orsay virus infection in *C. elegans* (**Fig 1B, C**). However, we found that Orsay virus infection does not modulate autophagic flux (**Fig 2**). In addition, re-feeding after starvation limits Orsay virus infection and blocks autophagic flux (**Fig 3**). Interestingly, we found that *atg-3* mutants phenocopy *rde-1* mutants in the transcriptional response to infection though they do not have a defect in RNAi (**Fig 4, 6**). In addition, we found that ATG-3 limits viral infection at a post-entry step, similar to RDE1 (**Fig 5**). Differential expression analysis upon bulk RNA sequencing revealed that *sqt-2*, which encodes a collagen trimer protein known to be important for Orsay virus control, is depleted in the context of infection as well as in naïve *atg-3* mutants (**Fig 7**), supporting regulation of collagen organization as a mechanism for ATG-3 limitation of Orsay virus replication.

Although we identified a novel role for ATG-3 in regulating Orsay virus infection, ATG-3 has several known functions in response to pathogens in different eukaryotic organisms. In mice, ATG3 plays a crucial role in the control of the parasite *Toxoplasma gondii* due to its involvement in targeting of LC3 and interferon-γ effectors onto the vacuolar membrane of *T. gondii* and its subsequent disruption (16). This effect has also been observed in control of bacterial pathogen *Chlamydia trachomatis*, which also replicates in pathogen-containing vacuoles (53). In addition, *Mycobacterium tuberculosis*, a bacterial pathogen, inhibits translation of ATG3 to promote pathogen survival, suggesting that ATG3 is relevant to defense against infection (54). ATG3 has been identified as part of the innate immune response in multiple agriculturally relevant plant pathogens such as tobacco mosaic virus (55, 56). Although *C. elegans* do not have IFNs, it is possible that the relationship between ATG-3 and structural components of defense against various pathogens are conserved in other eukaryotic organisms (31). Further characterization of the mechanism by which ATG-3 affects Orsay virus pathogenesis could thus have implications for treatment modalities for human-relevant pathogens.

We found that the viral susceptibility phenotype of *atg-3-*mutant *C. elegans* likely does not depend on the role of *atg-3* in maintaining autophagic flux. Indeed, if autophagic flux controlled Orsay virus infection, we would expect that disruption of other autophagy genes that affect autophagic flux such as *syx-17*, *epg-5*, and *rab-7* would also affect Orsay virus RNA levels. However, we observed that mutation of these genes does not change viral susceptibility compared to WT animals (**Fig 1D**). In addition, re-feeding after starvation blocked autophagic flux and reduced susceptibility to Orsay virus compared to unstarved animals, which further supports the idea that reduced autophagic flux does not cause increased susceptibility to Orsay virus infection (**Fig 3**).

We show for the first time with autophagic flux reporter animals that re-feeding after starvation suppresses autophagy pathways, likely due to the sudden change in nutrient availability, though it is well known that starvation induces autophagy. The status of nutrients such as zinc is known to affect both Orsay virus pathogenesis, and it is also relevant to the severity of many different human viral infections including SARS-CoV-2 (47, 57). Our finding that the history of nutrient status in *C. elegans* affects Orsay virus susceptibility is relevant to the standard protocol used in this model of viral infection, and also supports use of this model to more broadly investigate the role of nutrient status in viral infection. Although we ruled out neutral lipid accumulation as the mechanism by which Orsay virus infection is regulated in autophagy-mutant *C. elegans*, lipid metabolism pathways are complex, and several genes related to lipid metabolism including *sbp-1*, *fat-6/fat-7*, and *elo-5/elo-6* have been identified to be relevant to Orsay virus infection (47). It is possible that further study of lipid metabolism pathways affected by autophagy mutation could reveal a role for lipid metabolism in Orsay virus infection in *atg-3* mutants.

Our RNAseq findings were consistent with a previously published list of genes that are differentially expressed upon Orsay virus infection (42). Interestingly, we found that expression of *sqt-2*, which encodes a collagen trimer, is depleted upon Orsay virus infection of WT worms (**Fig 7A**). It is also depleted in naïve *atg-3* mutant *C. elegans* compared to naïve WT *C. elegans* (**Fig 7C**). SQT-2 has been recently identified as an antiviral factor that is expressed in the intestine in Orsay virus infection—intestine-specific RNAi knockdown of *sqt-2* led to increased viral susceptibility (31). We hypothesize that ATG-3 is involved in promoting the expression and organization of cellular collagens that have an antiviral role in Orsay virus infection, and the disruption of this pathway leads to the increased viral susceptibility of *atg-3-*mutant *C. elegans* (**Fig 1C**).

Infection by Orsay virus is known to cause structural changes within intestinal cells such as reorganization of the terminal web, diminished intermediate filaments, and liquefaction of the cytoplasm (18, 22). Is it possible that some of these cellular changes are mediated by interactions between Orsay virus and SQT-2. Though SQT-2 is thought to be part of the *C. elegans* cuticle, which forms the exoskeleton, its expression is relevant to Orsay virus infection in the intestine (31). SQT-2’s involvement in the extracellular matrix (ECM) could be relevant to viral entry or egress. Our data suggest that ATG-3 and its regulation of *sqt-2* could be more relevant to viral egress than viral entry, since bypassing viral entry in *atg-3-*mutant replicon worms led to a similar enhanced susceptibility to infection (**Fig 5**).

Another possibility is that *atg-3*-mutation leads to disruption of structural components on the apical side of the cell, which could include collagen encoded by *sqt-2* as well as other collagen and actin-related genes, preventing non-lytic egress of Orsay virus. The Orsay δ protein is required for non-lytic viral egress and is thought to interact with ACT-5, which is a component of the terminal web, to mediate egress (22). The accumulation of Orsay virus particles on the apical side of intestinal cells in *atg-3-*mutant *C. elegans* could be due to impaired viral egress rather than enhanced viral replication (**Fig 1I**). Further experiments to determine whether *atg-3* mutants shed Orsay virus at similar levels to *rde-1* mutants are needed to explore this possibility.

The relationship between autophagy, collagen, and the ECM is complex and requires further characterization. ATG-3 interactions with collagen are not well established in the literature, though there are suggestions that it may be conserved in vertebrates; in mice, expression of various collagen genes used as markers of fibrosis in the liver was reduced upon the inhibition of ATG-3 (58). Autophagy is also activated in response to damage of the *C. elegans* cuticle (59). The relationship between autophagy and the ECM is relevant to the progression of several different tumor types, as changes in the ECM have distinct pro- and anti-autophagic functions (60). In addition, type IV collagen has been identified as an important regulator of autophagic function in human muscle cells and neurons, and the regulation of autophagic function by type IV collagen is relevant to the progression of several different types of muscular dystrophy in humans (61, 62). Further exploration of the relationship between autophagy and collagen pathways in *C. elegans* could lead to important advances in our understanding of structural mechanisms of viral regulation within cells.

## METHODS

### *C. elegans* culture and maintenance

C. elegans VC2010, atg-13(bp414) III (HZ1688), epg-8(bp251) I; him-5(e1490) V (HZ1691), atg-9(bp564); him-5(e1490) V (HZ1687), atg-3(bp412) IV; him-5(e1490) V (HZ1684), syx-17(tm3182) (FX03182), epg-5(vk3748), rab-7(ok511)/mIn1 [mIs14 dpy-10(e128)] II (VC308), rict-1(ft7) II (KQ1366), and autophagic flux reporter worms were generously provided by Dr. Stephen Pak’s group at Washington University in St. Louis (40). C. elegans N2, jyIs8[Ppals-5::GFP;Pmyo-2::mCherry]; rde-1(ne219) V, and replicon worms were generously provided by Dr. David Wang’s group at Washington University in St. Louis. A full list of strains used can be found in **Table 1**. These strains were maintained under standard lab culture conditions (63). Animals were fed OP50 Escherichia coli (E. coli) on nematode growth medium (NGM) dishes stored at 20°C and passaged every 3 days to a fresh NGM dish seeded with OP50 E. coli.

**Table 1:** *C. elegans* lines.

| Name | Strain Name | Genotype | Reference | Source |
| --- | --- | --- | --- | --- |
| *atg-13* | HZ1688 | *atg-13(bp414) III* |  | CGC |
| *epg-8* | HZ1691 | *epg-8(bp251) I; him-5(e1490) V* |  | CGC |
| *atg-9* | HZ1687 | *atg-9(bp564); him-5(e1490) V* |  | CGC |
| *atg-3* | HZ1684 | *atg-3(bp412) IV; him-5(e1490) V (HZ1684)* |  | CGC |
| *syx-17* | FX03182 | *syx-17(tm3182)* |  | NBRP |
| *epg-5* | VK3763 | *epg-5(vk3748)* |  | Pak lab |
| *rab-7* | VC308 | *rab-7(ok511)/mIn1 [mls14 dpy-10(e128)]* |  | CGC |
| *rict-1* | KQ1366 | *rict-1(ft7) II* |  | CGC |
| *rde-1* | jyls8;rde-1(ne219) | {jyls8[Ppals-5::GFP;Pmyo-2::mCherry]; rde-1(ne219) V} | (27) | Wang lab |
| WT+ PHIP::RNA1 WT | N2;virEx64[PHIP::RNA1WT-1] | {virEx64[PHIP::OrsayRNA1 WT-1; Pmyo-2::YFP]} | (47) | Wang Lab |
| WT + PHIP::RNA1 D601A | N2;virEx66[PHIP::RNA1D601A-1] | {virEx66[PHIP::OrsayRNA1 D601A-1; Pmyo-2::YFP]} | (47) | Wang lab |
| *rde-1* + PHIP::RNA1 WT | jyls8;rde-1(ne219); virEx20[PHIP::RNA1WT-1] | {virEx20[PHIP::OrsayRNA1 WT-1; Pmyo-2::YFP]; jyls8[Ppals-5::GFP;Pmyo2::mCherry];rde-1(ne219) V} | (66) | Wang lab |
| *rde-1* + PHIP::RNA1 D601A | jyls8;rde-1(ne219); virEx21[PHIP::RNA1D601A-1] | {virEx21[PHIP::OrsayRNA1 D601A-1; Pmyo-2::YFP]; jyls8[Ppals5::GFP;Pmyo-2::mCherry];rde-1(ne219) V} | (66) | Wang lab |

CGC, *C. elegans* Genetics Center (www.cgc.umn.edu);
NBRP, National BioResource Project (www.nbrp.jp/en/resource/c-elegans-en)

### Orsay virus preparation, infection, and RNA extraction

Orsay virus was prepared as previously described (21). For all infection experiments, unless otherwise indicated, animals were bleached and then synchronized in M9 buffer in 15-mL conical tubes with constant rotation at room temperature for 18h. 200 arrested larval-stage 1 (L1) larvae were seeded into 6 well NGM plates seeded with OP50. L1 larvae were allowed to recover for 24h at 20°C prior to infection. Orsay virus filtrate was thawed at room temperature and then diluted 1:10 with M9 buffer. In each well, 20 μL of 2.5×10^5^ tissue culture infective dose (TCID_50_)/mL (multiplicity of infection (MOI) of 10) of virus filtrate was added to the bacterial lawn. The samples were incubated at and incubated at 20°C until sample collection. Animals were collected into 1.5-mL Eppendorf tubes by washing each well with 1 mL of M9 buffer either 48 hours post infection (hpi) or 72 hpi as indicated. Samples were pelleted by spinning for 1 min at 376 x *g* in a benchtop centrifuge. The supernatant was removed, and 300 μL of TRI reagent (Zymo Research Cat. No. R2050-1-200) was added to the tubes, which were then frozen in liquid nitrogen. For each experiment, six replicate wells were used for each infection condition unless otherwise indicated. Total RNA from infected animals was extracted using Direct-zol RNA miniprep (Zymo Research, Cat. No. R2053-A) purification according to the manufacturer’s protocol and eluted in 30 uL of RNase/ DNase-free water (47).

### Orsay virus quantification by RT-qPCR

One-step real-time quantitative reverse-transcription-PCR (RT-qPCR) was performed on RNA extracted from infected animals to quantify Orsay virus RNA as previously described (46). Briefly, RNA was diluted 1:100, and 5 μL was used in a TaqMan fast virus one-step qRT-PCR reaction (Applied Biosystems, Cat. No. 4444432) with primers and a probe (OrV_RNA2) that target the Orsay virus RNA2 segment. To control for variation in the number of animals, Orsay virus RNA levels were normalized to an internal control gene, *rps-20*, which encodes a small ribosomal subunit S20 protein required for translation in *C. elegans* (64). Data was analyzed with the ΔΔC_T_ method with the control group normalized to 1.0 in each individual experiment. At least two independent experiments were performed with six replicates each in each experiment.

### Transmission electron microscopy

For morphological analyses at the ultrastructural level, worms were fixed in 2% paraformaldehyde/2.5% glutaraldehyde (Ted Pella Inc., Redding, CA) in 100 mM cacodylate buffer, pH 7.2 for 2 hours at room temperature and then 2 days at 4°C. Samples were washed in cacodylate buffer and postfixed in 1% osmium tetroxide (Ted Pella Inc.)/ 1.5% potassium ferricyanide (Sigma, St. Louis, MO) for 1 hr. Samples were then rinsed extensively in dH20 prior to en bloc staining with 1% aqueous uranyl acetate (Ted Pella Inc.) for 1 hr. Following several rinses in dH20, samples were dehydrated in a graded series of ethanol and embedded in Eponate 12 resin (Ted Pella Inc.). Ultrathin sections of 95 nm were cut with a Leica Ultracut UCT ultramicrotome (Leica Microsystems Inc., Bannockburn, IL), stained with uranyl acetate and lead citrate, and viewed on a JEOL 1200 EX transmission electron microscope (JEOL USA Inc., Peabody, MA) equipped with an AMT 8 megapixel digital camera and AMT Image Capture Engine V602 software (Advanced Microscopy Techniques, Woburn, MA). Measurements of ribosomes and virus were made using the AMT software.

### Quantification of Autophagic Flux

Animals carrying the AFR were age-synchronized by seeding 20 adults, allowing them to lay eggs for 2 hours, and picking them off, yielding about 200 animals per plate. 25 adult animals at either 72h or 96h of life were placed in individual wells of 384-well plates, and images were automatically acquired and quantified using a plate reader type, high-content imager (Thermofisher CellInsight CX-7) as previously described (40). Three independent experiments were performed with four experimental replicates with four technical replicates each. GFP:mKate2 ratio was normalized to 1.0 in the control group in each individual experiment.

### Generation of atg-3 lines expressing Orsay RNA1 wild-type and RNA1 D601A mutant segments

We crossed the C. elegans WT + PHIP::RNA1 WT and WT + PHIP::D601A mutant lines into the *atg-3(bp412)* genetic background. To induce expression of the RNA1 segments, 20 adults stably expressing the transgenic RNA1 array were allowed to lay eggs for 2 hours at RT and then picked off the plate. The progeny were heat-shocked at 33°C for 2h as Day 3 adults. 24h after heat-shock, animals that express the RNA1 segment evidenced by the GFP head reporter were picked into M9 buffer, and fluorescence microscopy was performed.

### *In vivo* neutral lipid staining

LipidTox Red Neutral lipid (1000x, ThermoFisher H34476) was diluted in OP50 *E. coli* and seeded on NGM agar dishes to yield a final concentration of 10x. Animals were synchronized to the L1 stage as described above and transferred to the dish and incubated for 72 hours. Animals were washed off of plates with 1 mL M9 buffer and collected in 1.5 mL Eppendorf tubes. Samples were then washed 2x with M9 buffer after adult animals settled at the bottom of the tube. Microscopic imaging was performed as detailed below.

### Fluorescence Microscopy and Quantification

Animals were anesthetized with 25 mL tetramisole and then pipetted onto a 2% agarose pad and coverslipped. Images were acquired using an ApoTome microscope (Zeiss) and images were captured using Axiovision 4.8.2 software (Zeiss). Fluorescence was quantified using ImageJ software, and arbitrary fluorescence units were normalized to 1.0 in the control group.

### RNAi feeding for knockdown

RNAi feeding was used for gene knockdown as previously described (65). *E. coli* strain HT115 carrying double-stranded RNA expression cassettes for genes of interest was induced using established conditions and then seeded into NGM dishes containing IPTG. *pos-*1, *unc-22*, and *sqt-2* HT115 strains were from the Ahringer library (65).

### RNAseq

Total RNA integrity was determined using Agilent Bioanalyzer or 4200 Tapestation. Library preparation was performed with 1 to 3ug of total RNA. Ribosomal RNA was removed by a hybridization method using Ribo-ZERO kits (Illumina-EpiCentre). mRNA was then fragmented in reverse transcriptase buffer and heating to 94 degrees for 8 minutes. mRNA was reverse transcribed to yield cDNA using SuperScript III RT enzyme (Life Technologies, per manufacturer’s instructions) and random hexamers. A second strand reaction was performed to yield ds-cDNA. cDNA was blunt ended, had an A base added to the 3’ ends, and then had Illumina sequencing adapters ligated to the ends. Ligated fragments were then amplified for 12-15 cycles using primers incorporating unique dual index tags. Fragments were sequenced on an Illumina NovaSeq X Plus using paired end reads extending 150 bases. Basecalls and demultiplexing were performed with Illumina’s bcl2fastq software with a maximum of one mismatch in the indexing read.

RNA-seq reads were then aligned to the Ensembl release 101 primary assembly with STAR version 2.7.11a. (Two segments of Orsay virus (refseq: <u>NC_028097.1</u>, <u>NC_028098.1</u>) were manually added into the Ensembl release 101 databases with Cellranger (v. 7.2.0) pipeline for supplemental analysis).

All gene counts were then imported into the R and differential expression analysis was then performed with package DESeq2 (v. 1.34.0) for differences between conditions and the results were filtered for only those genes with Benjamini-Hochberg false-discovery rate adjusted p-values less than or equal to 0.05. Log2 transformed count table generated from DESeq2 was used for PCoA plot.

For each contrast extracted with DESeq, pathways were detected using the model organism database for C. elegansR Wormbase with R package enrich (v. 3.2), to test for changes in expression of the reported log 2 fold-changes reported by DESeq2 in each term versus the background log 2 fold-changes of all genes found outside the respective term.

Three sets of genes, representing genes induced by Orsay virus infection found by RNAseq, genes induced by Orsay virus found by RNAi screen, and a complete list of collagen genes were added into enrichR wormbase database to form three new pathways to accommodate our study design (31, 42, 51).

RNA-seq data was deposited at the European Nucleotide Archive (accession no. PRJEB74479).

### Statistical analysis

Statistical significance was determined with the Mann-Whitney test to compare two different groups of samples, and the Kruskal-Wallis test was used with *post hoc* comparisons analyzed by Dunn’s multiple comparison test when three or more samples were compared. A Spearman correlation test was used to analyze lipid/viral level correlations. Graphical representation and statistical analysis were performed using GraphPad Prism 9.

## Acknowledgements and funding

We acknowledge all members of the Baldridge laboratory for helpful discussions. We also acknowledge Dr. David Wang (Washington University School of Medicine) for provision of *C. elegans* reagents and critical advice. We thank Wandy Beatty and the Molecular Microbiology Imaging Facility for assistance in obtaining TEM images. We also thank the Genome Technology Access Center at the McDonnell Genome Institute at Washington University School of Medicine for assistance with genomic analysis. The Center is partially supported by NCI Cancer Center Support Grant #P30 CA91842 to the Siteman Cancer Center. This publication is solely the responsibility of the authors and does not necessarily represent the official view of NCRR or NIH.

G.K. was supported by National Institutes of Health (NIH) T32 AI007163 and T32 GM007200. M.T.B. was supported by NIH R01 AI173360 and an Investigator in the Pathogenesis of Infectious Disease award from the Burroughs Wellcome Fund. TS was supported by R35 GM152192. The funders did not play any role in the study design, data collection and analysis, decision to publish, or preparation of the manuscript.

## Notes

### Competing Interest Statement

The authors have declared no competing interest.

